# Extracellular Vesicles from Inflammation-Primed Adipose-Derived Stem Cells Enhance Achilles Tendon Repair by Reducing Inflammation and Promoting Intrinsic Healing

**DOI:** 10.1101/2023.01.31.526532

**Authors:** Hua Shen, Ryan A. Lane

## Abstract

Achilles tendon rupture is a common sports-related tendon injury. Even with advanced clinical treatments, many patients suffer from long-term pain and reduced function. These unsatisfactory outcomes result primarily from an imbalanced injury response with excessive inflammation and inadequate regeneration. Prior studies showed that extracellular vesicles from inflammation-primed adipose-derived stem cells (iEVs) can attenuate inflammation in the early phase of tendon healing. However, the effect of iEVs on tendon inflammation and regeneration in the later phases of tendon healing and the underlying mechanism remain to be determined. Accordingly, this study investigated the mechanistic roles of iEVs in regulating tendon response to injury using a mouse Achilles tendon injury and repair model in vivo and iEV-macrophage and iEV-tendon cell co-culture models in vitro. Results showed that iEVs promoted tendon anti-inflammatory gene expression and reduced mononuclear cell infiltration in the remodeling phase of tendon healing. iEVs also increased injury site collagen deposition and promoted tendon structural recovery. As such, mice treated with iEVs showed less peritendinous scar formation, much lower incidence of postoperative tendon gap or rupture, and faster functional recovery compared to untreated mice. Further in vitro study revealed that iEVs both inhibited macrophage inflammatory response and increased tendon cell proliferation and collagen production. The iEV effects were partially mediated by miR-147-3p, which blocks the toll-like receptor 4/NF-κB signaling pathway that activates macrophage M1 polarization. The combined results demonstrated that iEVs are a promising therapeutic agent, which can enhance tendon repair by attenuating inflammation and promoting intrinsic healing.

**Significance statement:** Using a clinically relevant mouse Achilles tendon injury and repair model, this study revealed that iEVs, a biological product generated from inflammation-primed adipose-derived stem cells, can directly target both macrophages and tendon cells and enhance tendon structural and functional recovery by limiting inflammation and promoting intrinsic healing. Results further identified miR-147-3p as one of the active components of iEVs that modulate macrophage inflammatory response by inhibiting toll-like receptor 4/NF-κB signaling pathway. These promising findings paved the road toward clinical application of iEVs in the treatment of tendon injury and other related disorders.

## Introduction

Tendon injury, accounting for nearly half of all sports-related injuries, is one of the most common and challenging orthopedic conditions.^1–5^ Being the largest and strongest tendon in the body, the Achilles tendon is one of the most injured tendons due to sports, exercise, or athletic activities.^4,6^ The tendon attaches gastrocnemius and soleus muscles to the calcaneus bone. By transmitting muscle force to the calcaneus, the Achilles tendon enables plantar flexion of the foot, which is required for varied types of locomotion, such as walking, running, and jumping. Current clinical interventions after Achilles tendon rupture include surgical or non-surgical treatments and rehabilitation.^7^ Despite state-of-the-art surgical techniques and rehabilitation methods, many patients suffer from long-term pain and reduced function following Achilles tendon rupture. A clinical study in a general population reported that major functional deficits persist 2 years after acute Achilles tendon rupture.^8^ Others reported that calf muscle performance deficits remain 7 years after an Achilles tendon rupture.^9^ Similarly, 30%–40% of professional players were unable to return to sports after Achilles tendon ruptures; those who did return played in fewer games, had less play time, and performed at a lower level than before the injury,^10,11^ Thus, there is a great need for new treatments to effectively improve Achilles tendon healing after injury.

The unsatisfactory outcomes after tendon injury are associated with excessive inflammation mainly driven by infiltrating macrophages and inadequate regeneration mediated by resident tendon cells, including tenocytes, tendon stem/progenitor cells, and epitenon cells.^5,12–17^ Activated macrophages primarily exhibit two functional phenotypes: a default pro-inflammatory M1 phenotype and an alternative anti-inflammatory and pro-regenerative M2 phenotype.^18^ Although inflammation is required to initiate healing process and clear damaged tissues, excessive/sustained inflammation causing tendon cell death, matrix degradation, and peritendinous scar formation impedes tendon structural and strength recovery and excursion.^12,13,19–21^ Moreover, the collagen-producing tendon cells lack the capacity to repair injured tissue intrinsically.^5,14–17^ Therefore, our treatment strategy has focused on limiting macrophage inflammatory response and stimulating tendon cell activity and function to promote scarless tendon healing and functional recovery.

We recently found that extracellular vesicles generated by inflammation-primed adipose-derived stem cells (iEVs) can reduce inflammation in the early phase of tendon healing.^22^ It remains to be determined if iEVs may also promote tendon structural and functional recovery in the later phases of tendon healing and the underlying cellular and molecular mechanisms. Extracellular vesicles (EVs) are nanosized vesicles carrying varied types of active molecules including microRNAs (miRNAs) and mRNAs that mediate EV functions.^23,24^ While nearly all types of cells can generate EVs, EVs from different types of cells and cells at different functional states carry different cargo molecules and thereby exbibit distinct therapeutic potential.^22,25–28^ We hypothesized that iEVs from inflammation-primed adipose-derived stem cells (iASCs) can both reduce inflammation and stimulate intrinsic tendon healing by shuttling active molecules that regulate macrophage and tendon cell functions. To test the hypothesis, this study investigated the dose-effect of iEVs on tendon inflammatory and healing responses up to 28 days after Achilles tendon injury and repair in a preclinical mouse model. The mechanisms of iEV actions including active cargo molecules were explored in vitro using iEV-macrophage and iEV-tendon cell co-culture models. Our results show that iEVs are a promising therapeutic agent for tendon repair.

## Materials and Methods

### Animals

Adult NF-κB-GFP-luciferase transgenic reporter mice (NGL; Jackson Laboratory; 4-5 months old) of both sexes were used for isolating adipose-derived stem cells (ASCs), preparing bone marrow-derived macrophages, and conducting all *in vivo* studies. ScxGFP tendon reporter mice (2-3 months old of both sexes) were used for tendon cell isolation.^29^ All experimental procedures were conducted in accordance with the Public Health Service Policy on Humane Care and Use of Laboratory Animals and approved and overseen by the Washington University Institutional Animal Care and Use Committee.

### Cell isolation and culture

ASCs expressing the mesenchymal stem cell markers CD29, CD44, and CD90 were isolated from subcutaneous fat of adult NGL mice and cultured in Minimum Essential Medium Alpha (alpha-MEM; Mediatech, Inc.) containing 10% fetal bovine serum (FBS; Life Technologies) as previously described.^22,30^ Macrophages were prepared from bone marrow monocytes of adult NGL mice and cultured in a macrophage culture medium containing 10% conditioned medium from L929 cell culture, 100 unit/ml penicillin, 100 μg/ml streptomycin (Life Technologies), and 10% FBS in alpha-MEM as detailed elsewhere.^22^ Tendon cells were isolated from tail tendons of adult ScxGFP mice and cultured in Dulbecco’s Modified Eagle’s Medium (DMEM, Mediatech, Inc.) supplemented with 10% FBS.^27^ Over 90% of isolated tendon cells expressed ScxGFP and about 0.3% ScxGFP+ cells co-expressed CD146.

### Preparation of extracellular vesicles

Naïve EVs and iEVs were isolated from untreated naïve ASCs and iASCs, respectively. iASCs were prepared by pretreating ASCs with interferon γ (100 ng/ml; R&D Systems) in ASC culture medium for 24 hours. After being washed with Dulbecco’s phosphate buffered saline (DPBS; Life Technologies) for three times, both naïve ASCs and iASCs were cultured in alpha-MEM containing 2% EV-free FBS for 48 hours. The resulting conditioned medium was cleared via centrifugation at 500×g for 10 min and 10,000×g for 30 min at 4°C. Vesicles in the cleared medium were collected by ultracentrifugation at 100,000×g for 70 min at 4°C and resuspended in DPBS as previously described.^22^

### Characterization of extracellular vesicles

Vesicle morphology was examined via transmission electron microscopy (TEM).^22^ Vesicle size and concentration were determined by Nanoparticle Tracking Analysis (NTA; NanoSight NS300, Malvern Instruments Ltd.). CD9+/CD63+/CD81+ iEVs were separated from bulk iEVs with Exo-Flow streptavidin magnetic microbeads (System Biosciences) coupled with biotinylated human anti-mouse CD63 (Miltenyi 130-125-989), rat anti-mouse CD9 (Miltenyi 130-101-961), and hamster anti-mouse CD81 (Miltenyi 130-101-966) antibodies according to manufacturer’s instructions. Half of iEVs captured by the beads were stained with Exo-FITC Universal Exosome Stain (System Biosciences) and analyzed with a BD FACSCanto™ II Flow Cytometry System, using microbeads prepared in parallel without capturing antibodies as negative controls. The remaining half of captured iEVs were eluted from microbeads with Exosome Elution Buffer (System Biosciences), washed with DPBS, and subjected to NTA analysis to determine vesicle size and percentage in total iEV population.

### Macrophage inflammatory response

Macrophages expressing the NGL NF-κB-luciferase reporter were either pre-treated with iEVs at a dose corresponding to an iEV donor and recipient cell ratio of 20:1^22^ or transfected with a mirVana miRNA mimic mmu-miR-147-3p (Thermo Fisher Scientific) with Lipofectamine™ RNAiMAX Transfection Reagent (Life technologies) in alpha-MEM containing 2% EV-free FBS and 10% EV-free conditioned medium from L929 cell culture for 48 hours. Cells pre-treated with equal volume of DPBS or control mimics in the same medium were used as iEV or miRNA mimic negative controls. The pretreated cells were stimulated with Lipopolysaccharides (LPS, 100ng/mL; Sigma-Aldrich) and IFNγ (50 ng/mL) in alpha-MEM containing 10% FBS and 10% L929 conditioned medium for 24 hours. The resulting conditioned medium was collected and assessed for inflammatory cytokine and chemokine concentrations using a ProcartaPlex multiplex immunoassay kit (Thermo Fisher Scientific). The cells were lysed and assessed for NF-κB activity with an Illumination Firefly Luciferase Enhanced Assay Kit (Gold Biotechnology). The results were normalized by samples’ protein concentrations.

### Tendon cell activity and function

Tendon cells were treated with iEVs at a dose corresponding to an iEV donor and recipient cell ratio of 20:1 or miR-147-3p mimics as above described in DMEM containing 2% EV-free FBS for 48 hours. Conditioned medium from the culture was collected to determine tendon cell type I collagen production using a Mouse Type I Collagen Detection Kit (Chondrex). The cells were lysed to assess cell proliferation with a CyQUANT™ Cell Proliferation Assay (Thermo Fisher Scientific) according to the manufacturer’s instructions. Type I collagen production were normalized by tendon cell counts.

### In vivo study design

Our previous study showed that iEVs from interferon γ-primed iASCs but not EVs from naïve ASCs have the potential to enhance Achilles tendon healing.^22^ Therefore, this study focused on the long-term dose effect of iEVs on tendon inflammatory and healing responses. 32 NGL mice (15 males and 17 females) from 4 litters were subjected to right Achilles tendon 2/3 transection and suture repair. Mice from two of the four litters (N=17) were pre-assigned for a video-based gait analysis to assess tendon functional recovery after injury.^31^ Mice from the other two litters (N=15) were pre-assigned for live bioluminescence imaging (BLI) to determine repair site NF-κB activity as described elsewhere.^22^ The repaired animals were divided into three litter- and gender-matched groups (N=10-11/group) and treated with 0 (CTRL), 1E+09 (+iEVL), and 5E+09 (+iEVH) iEVs loaded on a collagen sheet. BLI (N=5-6/group) was conducted 1 day before and 4, 7, 13, and 28 days after tendon repair. Function assessments (N=5-6/group) were performed 1-2 days before and 13, 20, and 27 days after tendon repair. All repaired mice were euthanized 28 days after injury and repair. The integrity of repaired tendons was first assessed under a dissecting microscope. Postoperative gap formation and rupture were defined as partial and complete loss of the continuity of the repaired tendons, respectively.^22^ Tendon inflammatory and healing responses were further assessed histologically (N=3-4/group) and quantitatively by TaqMan PCR for changes in tendon gene expression (N=7-8/group) using intact tendons from age and gender matched healthy NGL mice as intact tendon controls (N=8).

### Mouse Achilles tendon injury, repair, and treatment

The methods for mouse right Achilles tendon 2/3 transection, suture repair, and local delivery of iEVs via a collogen sheet were described previously.^12,22^ In brief, Achilles tendon transection was performed at the midpoint level between the calcaneal insertion and the musculotendinous junction under isoflurane anesthesia. The transected tendon was immediately repaired with a two-strand modified Kessler technique.^22^ iEVs were pre-loaded to the surface of one side of a collagen sheet containing 2 mg/ml type I collagen from rat tail (Corning Life Sciences) and applied around the surface of repair site with the iEV-loaded side facing repaired tendon.^12,22^ After recovery from anesthesia, repaired mice were returned to their home cages without physical restraint.

### Tendon histology

Mouse Achilles tendons attached to the calcaneus bone and gastrocnemius muscle were fixed with 4% paraformaldehyde in phosphate buffered saline (PBS), decalcified in 14% ethylenediaminetetraacetic acid (pH 7.2; Sigma-Aldrich), and embedded in paraffin. Serial sagittal sections (5μm thick) were then prepared and stained with a Russell Movat Pentachrome Stain Kit (StatLab). The section plane was defined by the presence of entire muscle-tendon-bone unit connected by the Achilles tendon including the calcaneal enthesis. The pentachrome stain reveals cell nuclei and elastic fibers in dark purple to black color, collagen in yellow, and intensely acidophilic collagen fibers in red on a yellow background.^22,32^ Stained sections were scanned with a NanoZoomer Whole Slide Imaging System (2.0-HT, Hamamatsu). Images at the injury site were exported at 40× magnification for cellularity and collagen content assessments and 10× magnification for scar tissue analysis. The cellularity of repaired tendons was determined by counting the number of fibroblast-like cells and mononuclear cells in the tendon region of exported images. The posterior and fat pad scar areas were defined as the area occupied by scar tissue posterior to repaired tendon and the area occupied by scar tissue at the fat pad region anterior to repaired tendon, respectively. The scar tissue area and collagen area were determined with the area analysis and color range selection tools of Adobe Photoshop CC 2015.5 (Adobe Systems Incorporated) as previously described.^22^ Every sample was assessed blindly on two different sections at the level approximately halfway from the tendon surface. The results were averaged and normalized by tissue size.

### Ankle joint angle measurement

Because the Achilles tendon is required for plantar flexion of ankle joint, the angle of ankle joint was used as a measurement of tendon functional recovery.^31^ The angle was determined when mice were at a full standing position (i.e., all paws are in contact with treadmill belt) while running on a rodent treadmill (EXER 3/6, Columbus Instruments) at a speed of 6~8 m/min. All mice were pretrained at the same speed for 10 minutes per day for 3 consecutive days. A 30-second video that recorded the sagittal view of mouse during running was taken with a digital camera on the indicated days before and after tendon injury and repair. All frames with mouse at the full standing position (11±2 frames per mouse per time point) in each video were analyzed with Image J 1.52a. for the ankle joint angle formed from the fibular head and the 5th metatarsal head to calcaneal tuberosity.

### RNA isolation and quantitative RT-PCR

Total RNA from cultured cells and mouse Achilles tendons were isolated with TRIzol reagent and RNeasy Mini Spin Column (Qiagen Sciences) and reversely transcribed into cDNAs using a SuperScript IV VILO Master Mix (Life Technologies).^22,30^ The relative abundances of genes of interest were determined by TaqMan PCR using primers and probes purchased from Applied Biosystems TaqMan Gene Expression Assays. *Ipo8* was used as an endogenous reference gene. Changes in tendon gene expression were determined by the comparative Ct method and expressed as relative mRNA abundance.

### EV miRNA TaqMan PCR

EV RNA was isolated with a SeraMir Exosome RNA Column Purification Kit (System Biosciences) according to manufacturer’s instructions. RNA integrity was determined by an Agilent 2100 bioanalyzer. cDNA of EV miRNA was synthesized from 10 ng total RNA using a TaqMan MicroRNA Reverse Transcription Kit (Applied Biosystems) and primers specific for mmu-miR-147-3p (alias, hsa-miR-147b; Applied Biosystems). The relative abundance of EV miRNA was determined by TaqMan PCR using TaqMan Universal PCR Master Mix II and TaqMan primers and probes from Applied Biosystems. hsa-miR-423-3p was used as an endogenous reference miRNA, as it was identified as one of the most abundant and stable miRNAs in ASC EVs by our small RNA-sequencing analysis.

### Small RNA-sequencing and data analysis

cDNA library preparation, RNA-sequencing, data acquisition, quality control, and processing were performed by the Washington University Genome Technology Access Center. In brief, 5μl of each RNA sample was used as input into a TruSeq Small RNA library Preparation kit (Illumina) per manufacturer’s instruction. cDNA libraries were amplified for 11 cycles with oligos adding a unique index. The resulting samples were sized and quantified with a Bioanalzyer High Sensitivity DNA Analysis kit (Agilent). A region table was used to determine molar concentration between 145–160bp, and an equimolar amount was pooled and then size selected per kit instruction. Libraries were run on HiSeq 3000 (Illumina) using single reads extending 50 base pairs. Differentially expressed miRNAs were identified using generalized linear models and filtered for False Discovery Rate (FDR) adjusted p-values < = 0.05.

### Statistical Analysis

Unless described elsewhere, all data are shown as mean ± standard deviation. Two-tailed paired t-tests and one-way analysis of variance (ANOVA) followed by Tukey’s or Dunn’s multiple comparisons tests (when appropriate) were used for two- and multiple-group comparisons of in vitro results. Two-way repeated measures ANOVA followed by FDR multiple comparisons tests with two-stage step-up procedure of Benjamini, Krieger and Yekutieli was used to assess the longitudinal impact of iEV treatment on tendon NF-κB activity. Two-way repeated measures ANOVA followed by Tukey’s multiple comparisons tests was used to evaluate the impact of iEVs on ankle joint angle recovery. One-way ANOVA followed by Tukey’s or Dunn’s multiple comparisons tests (when appropriate) was used to evaluate the effect of iEVs on tendon gene expression and tendon histomorphometry. Outliers were identified using the ROUT method with an FDR of 1%.^33^ All statistical analyses were performed with Prism 9 (GraphPad Software, LLC). Significance was set at p < 0.05.

## Results

### Characterization of iEVs

iEVs are pancake-like and of varied sizes as revealed by TEM imaging (pointed by white arrows in Figure 1A). NTA analysis detected a mean mode size of 146.4±12.4 nm for bulk iEVs (N=3 independent isolations; Figure 1B). Magnetic bead-assisted vesicle sorting followed by flow cytometry and NTA analyses showed that iEVs contained CD9+/CD63+/CD81+ exosomes (a major subtype of EVs; Figure 1C), which were smaller (mean mode size: 105.4±6.2 nm, N= 4 independent isolations) than bulk iEVs (p = 0.002) and accounted for about 70% of total iEVs.

**Figure 1.**
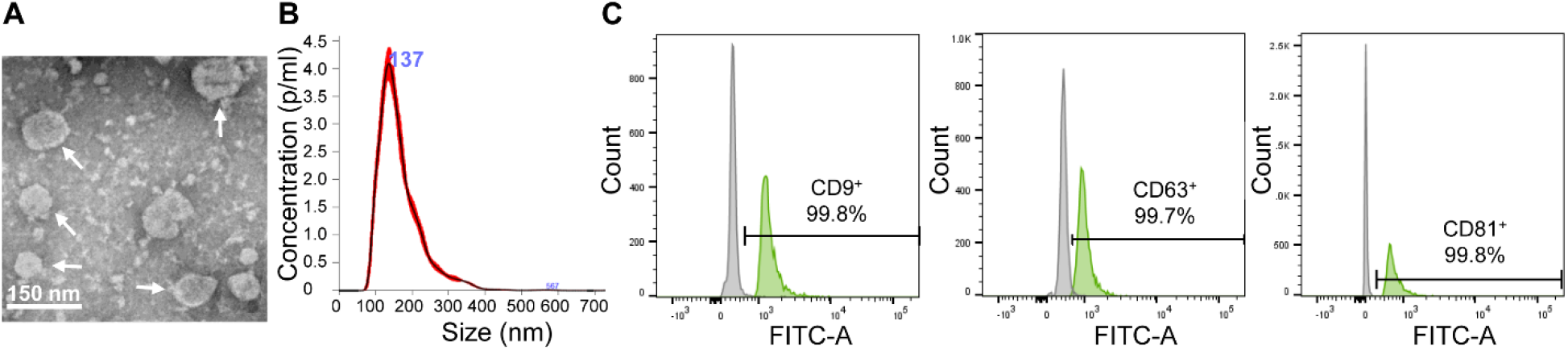
Characterization of extracellular vesicles from inflammation-primed adipose-derived stem cells (iEVs). **(A)** A representative transmission electron microscopy image of iEVs (indicated with white arrows); scale bar = 100 nm. **(B)** Representative nanopartical tracking analysis of iEVs. **(C)** Representative flow cytometry analysis of iEVs sorted with microbeads coupled with (green) or without (grey) biotinylated antibodies against indicated exosome markers.

### The dose-effect of iEVs on tendon inflammatory responses to acute injury and repair

Tendon inflammatory response after Achilles tendon injury and repair was first assessed longitudinally in live mice via bioluminescence imaging for injury site NF-κB activity. Results revealed an over 3-fold increase in NF-κB activity in untreated control mice (CTRL) 1 and 4 days after repair (Figure 2A). The increase was modestly reduced but remained significantly higher than the preinjury level 7 and 13 days after repair. iEV treatment at either dose effectively shortened the inflammatory response. Higher than preinjury level NF-κB activity was only detected in iEV-treated mice (+iEVL and +iEVH) 1 and 4 days after repair. High dose iEVs were more effective than low dose iEVs. The injury-site NF-κB activities in iEVH- but not iEVL-treated mice were much lower than those in control mice at all time points assessed within the first 2 weeks (i.e., 1, 4, 7 and 13 days) after tendon injury and repair (Figure 2A).

**Figure 2.**
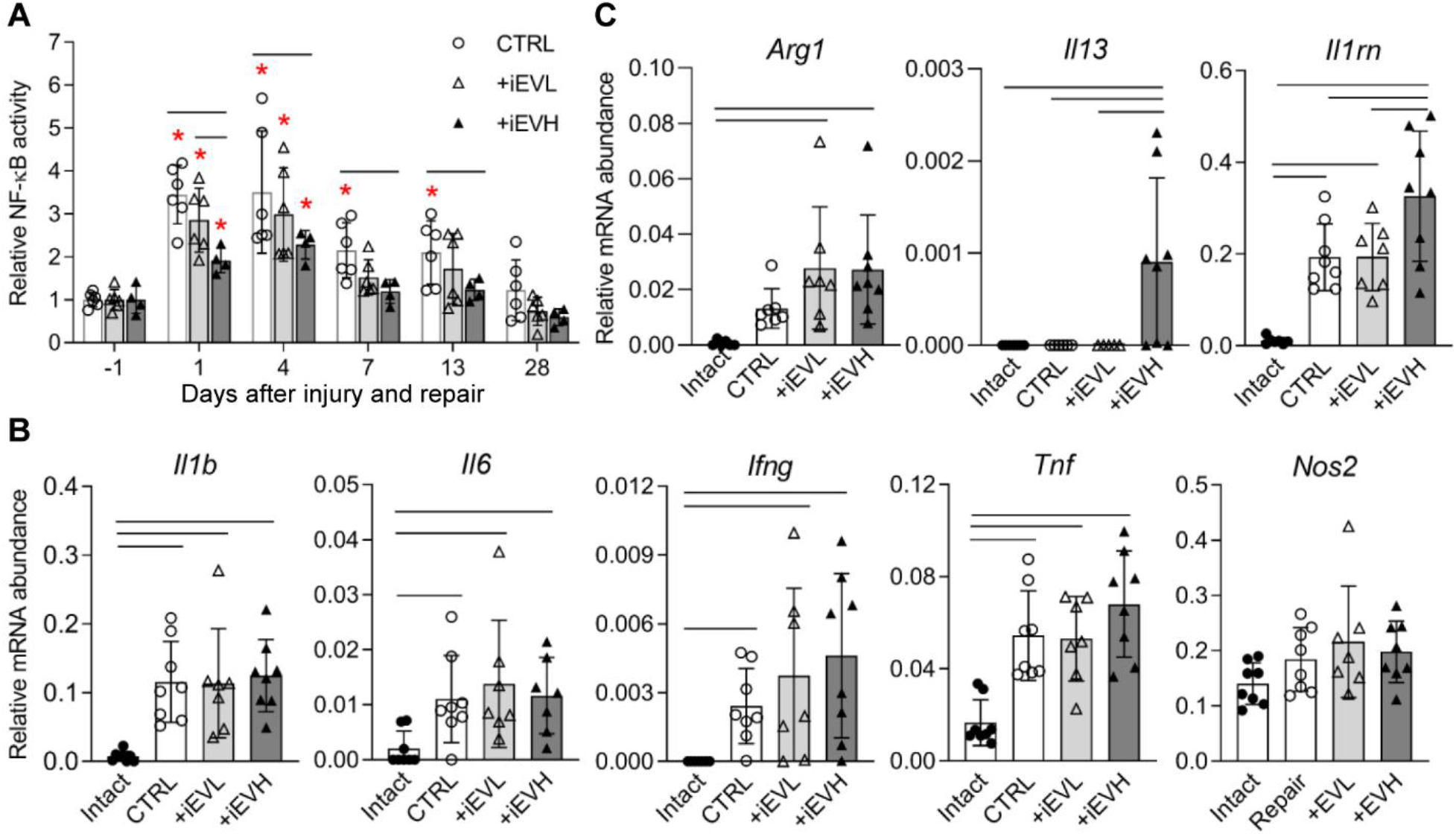
iEVs dose-dependently attenuate tendon inflammatory response to acute injury and repair. (**A**) Bioluminescence imaging revealed differential changes in injury site NF-κB activity in mice treated with 0, 1E+09, and 5E+09 iEVs (CRTL, +iEVL, and +iEVH) on the indicated days after Achilles tendon injury and repair. *, p < 0.05 compared to the pre-injury level (−1 day) of corresponding groups. —, p < 0.05 between indicated groups. (**B and C**) Differential expressions of inflammatory (B) and anti-inflammatory (C) genes in intact and repaired tendons subjected to the indicated treatments 28 days after injury and repair. —, p < 0.05 between indicated groups.

Gene expression analysis was subsequently performed in repaired tendons 28 days after injury and indicated treatments. Results showed that the expression levels of many but not all inflammatory genes (i.e., *Il1b, Il6, Ifng,* and *Tnf* but not *Nos2,* Figure 2B) were higher than normal, and no significant differences were detected between control and iEV-treated tendons. Despite these increases, the relative abundances of these genes in the remodeling phase were either essentially very low (*Il6* and *Ifng,* Figure 2B,) or substantially reduced to less than 5% (*Il1b,* Figure 2B, p < 0.001) and 35% (*Tnf,* Figure 2B, p <0.001) of what previously detected in untreated control tendons 7 days after injury and repair.^22^ Notably, tendons treated with iEV at either dose expressed higher-than-normal levels of *Arg1,* a marker for the anti-inflammatory M2 macrophages (Figure 2C). High dose iEVs also dramatically increased the expressions of anti-inflammatory genes *Il13* that activates macrophage M2 phenotype and *Il1rn,* an IL1β-inhibitor (Figure 2C), thus supporting a positive role of iEVs in promoting macrophage M2 polarization.

Histological assessment was conducted on pentachrome-stained tendon sections 28 days after injury and repair. Results revealed that tendon injury led to massive scar tissue formation along the posterior surface (red crosses in Figure 3A) and around the fat pad region of untreated tendons (white asterisk in Figure 3A) coupled with extensive mononuclear cell accumulation to the injury site (red arrows in Figure 3D). In accordance with their effects on the NF-κB activity and anti-inflammatory gene expression, iEV treatments at either dose effectively limited both scar tissue formation and mononuclear cell accumulation (Figure 3A-3I).

**Figure 3.**
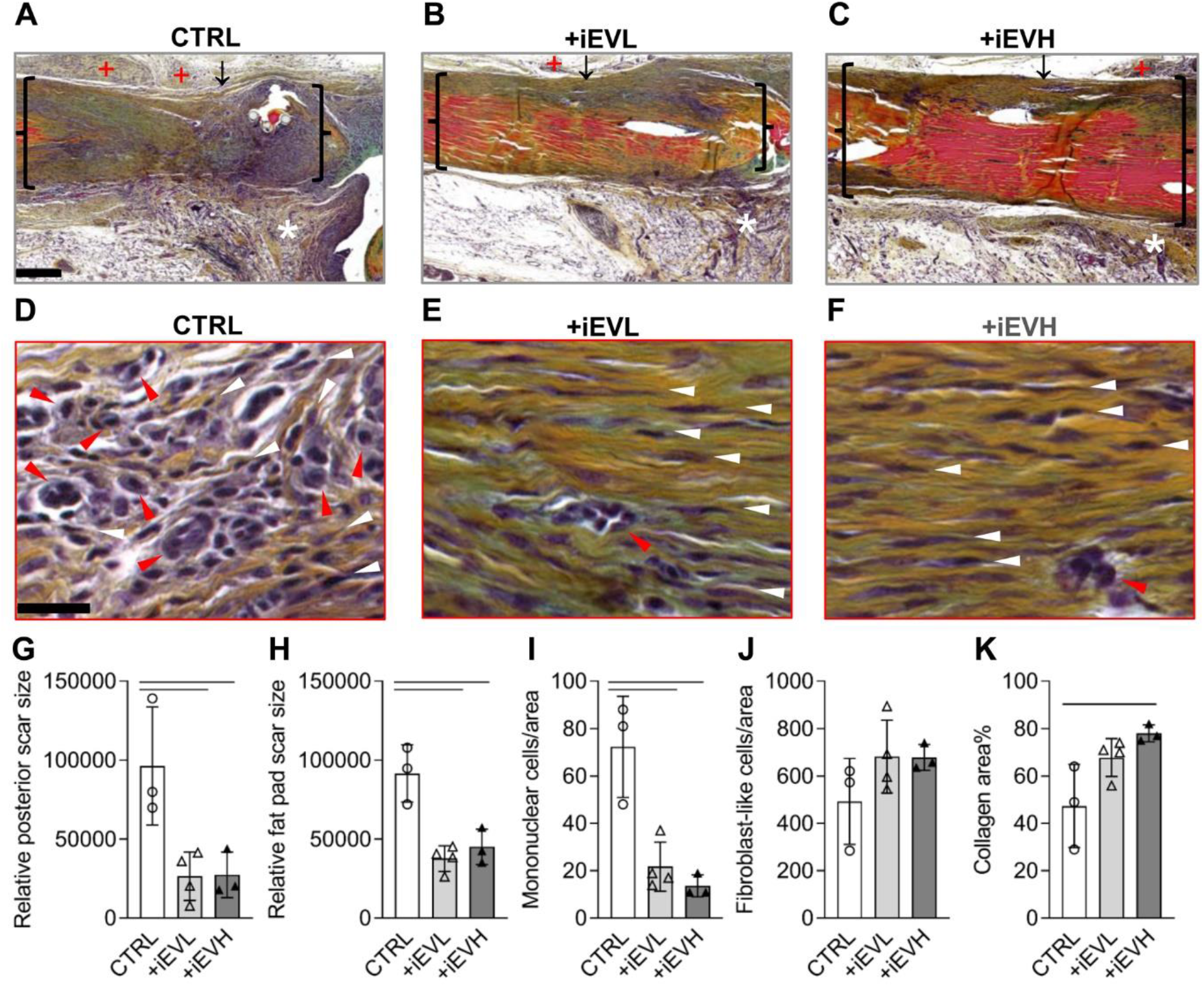
iEVs dose-dependently regulate Achilles tendon remodeling 28 days after injury and repair. **(A-F)** Representative images of pentachrome-stained Achilles tendon sections from mice treated with 0, 1E+09, or 5E+09 iEVs (CRTL, +iEVL, or +iEVH). Black braces and white asterisks in A-C mark the peripheral borders of repaired Achilles tendons and the fat pad region. ↓, transection site; +, posterior scar; Red and white arrowheads in D-F point to mononuclear cells and fibroblast-like cells, respectively. Scale bar in A is equal to 250μm and applies to A-C. Scale bar in D is equal to 25 μm and applies to D-F. **(G-K)** Semiquantitative assessments of the effect of iEVs on peritendinous scar formation (G and H), tendon cellularity (I and J), and collagen deposition (K). —, p < 0.05 between indicated groups.

### The dose-effect of iEVs on tendon remodeling after injury and repair

Changes in expression of tenogenic and tendon matrix genes were assessed in Achilles tendons 28 days after injury and repair. Results showed extensive increases in expression of all genes evaluated in untreated control tendons compared to intact tendons (Figure 4). Interestingly, tendons treated with either dose of iEVs showed less *Tnmd* increase compared to control tendons. High but not low dose iEVs also suppressed *Scx* and *Col1a1* increases in the remodeling phase (Figure 4).

**Figure 4.**
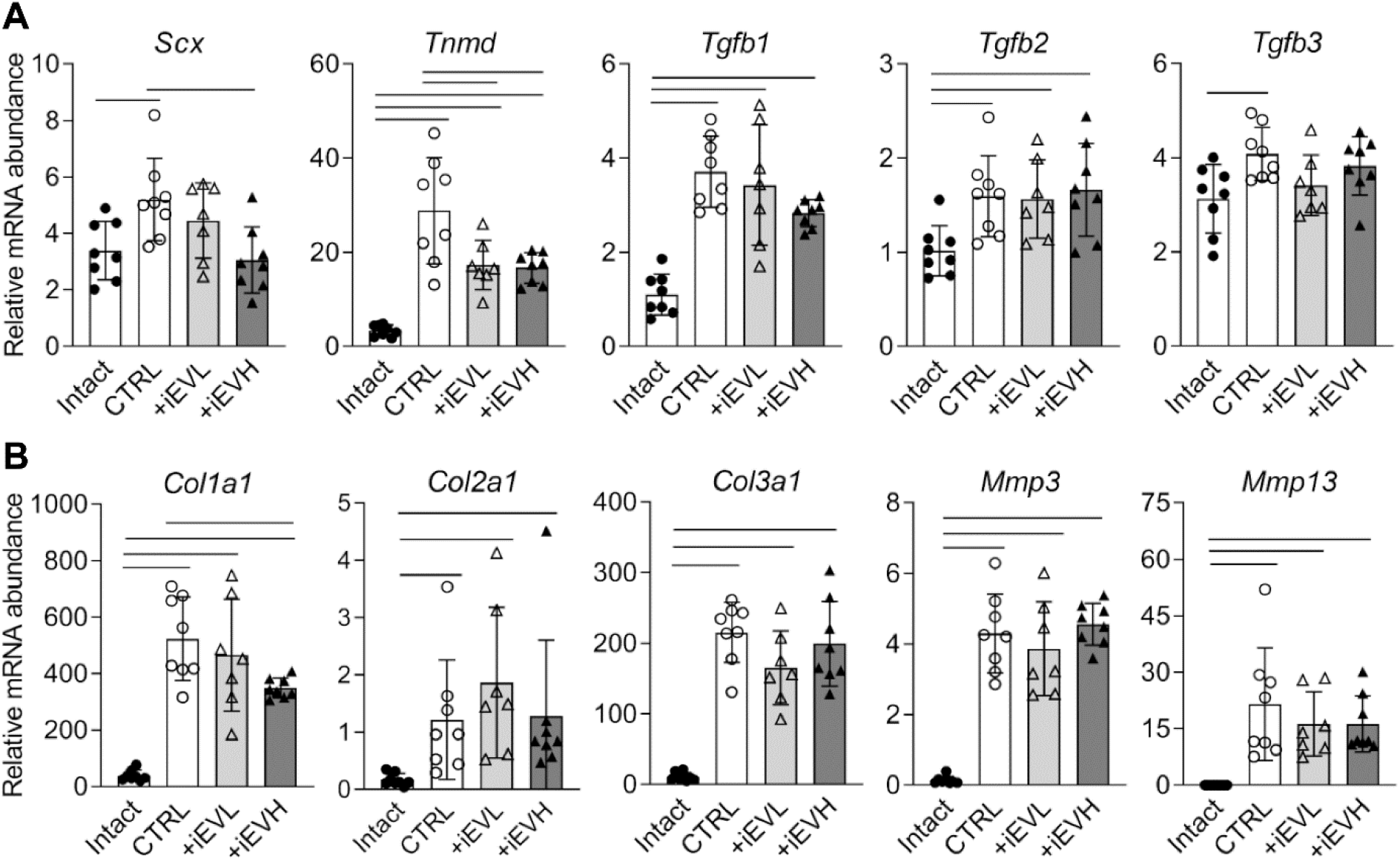
iEVs modulate tendon gene expression in the remodeling phase of tendon healing. **(A and B)** Quantitative PCR revealed differential expression of genes involved in tendon cell growth and differentiation (A) and matrix remodeling (B) in intact and repaired tendons treated with 0, 1E+09, or 5E+09 iEVs (CRTL, +iEVL, or +iEVH) 28 days after Achilles tendon injury and repair. —, p < 0.05 between indicated groups.

In line with increased tenogenic gene expression, histological assessment revealed robust fibroblast-like cell accumulation to the injury center of repaired tendons from all groups (white arrowheads in Figure 3D-3F). While there were no significant differences in fibroblast-like cell density among the three repair groups (Figure 3J), cells in iEV-treated tendons were better aligned with the long axis of Achilles tendon than those in untreated tendons (white arrowheads in Figure 3D-3F). With controlled *Col1a1* expression, high dose iEV both significantly increased collagen accretion in the injury center and effectively limited peritendinous scar formation after tendon injury and repair (Figure 3A-3F, 3G, 3H, and 3K), demonstrating a positive role of iEVs in balancing tendon injury response.

### The dose-effect of iEVs on tendon repair outcome

Mouse functional recovery after Achilles tendon injury and repair was assessed in live mice by a video-based gait analysis for ankle joint angle of repaired limbs. The angle of all mice assessed was slightly over 90° prior to injury and reduced to around 80° 13 days after injury and repair (Figure 5). In untreated mice, the defect was slowly reduced at a rate of about 2° per week but remained significantly lower than preinjury level by 27 days after repair. Treatment with low and high dose iEVs significantly increased the recovery rate by 30% and 43%, respectively. As a result, by 27 days after repair, untreated mice showed nearly 50% functional deficit compared to preinjury level. By contrast, iEVL- and iEVH-treated mice experienced only about 20% deficits. Tendon gap and rupture formation were examined postmortemly 28 days after injury and repair. In accordance with improved tendon structural and functional recovery, only one gap formation was noted in total 11 repairs treated with low dose iEVs. Likewise, one incidence of gap formation was detected in total 10 repairs treated with high dose iEVs. By contrast, 2 gap formations and 3 ruptures were found in total 11 untreated repairs.

**Figure 5.**
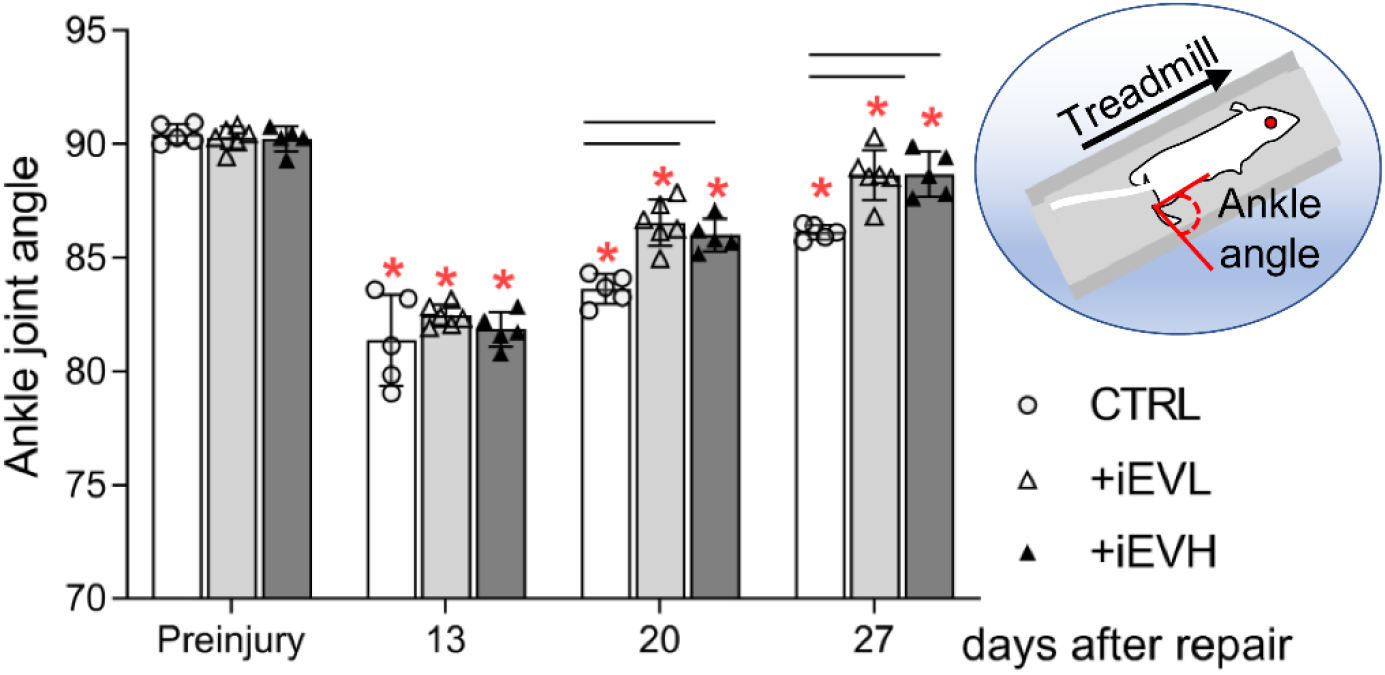
Changes in ankle joint angle in mice treated with 0, 1E+09, or 5E+09 iEVs (CRTL, +iEVL, or +iEVH) on the indicated days after Achilles tendon injury and repair. *, p < 0.05 compared to respective preinjury level. —, p < 0.05 between indicated groups.

### The in vitro effect of iEVs on macrophage inflammatory response

The combined in vivo findings from this and our prior study supported that iEVs attenuate tendon inflammatory response by inhibiting macrophage NF-κB signaling and promoting an M1 to M2 phenotypic switch.^22^ To explore this further, we stimulated bone marrow-derived macrophages with TLR4 agonists LPS and IFNγ, which activated NF-κB (Figure 6A), leading to an M1 phenotype with mass release of inflammatory cytokines IL-1β, TNFα, and IL-6 and chemokines CXCL1 and CXCL10 but not M2 phenotype stimuli IL-4 and IL-13 (Figure 6B). As expected, pretreatment of macrophages with iEVs inhibited NF-κB activation by LPS and IFNγ (Figure 6C) and dramatically reduced IL-1 and IL-6 release by macrophages (Figure 6D).

**Figure 6.**
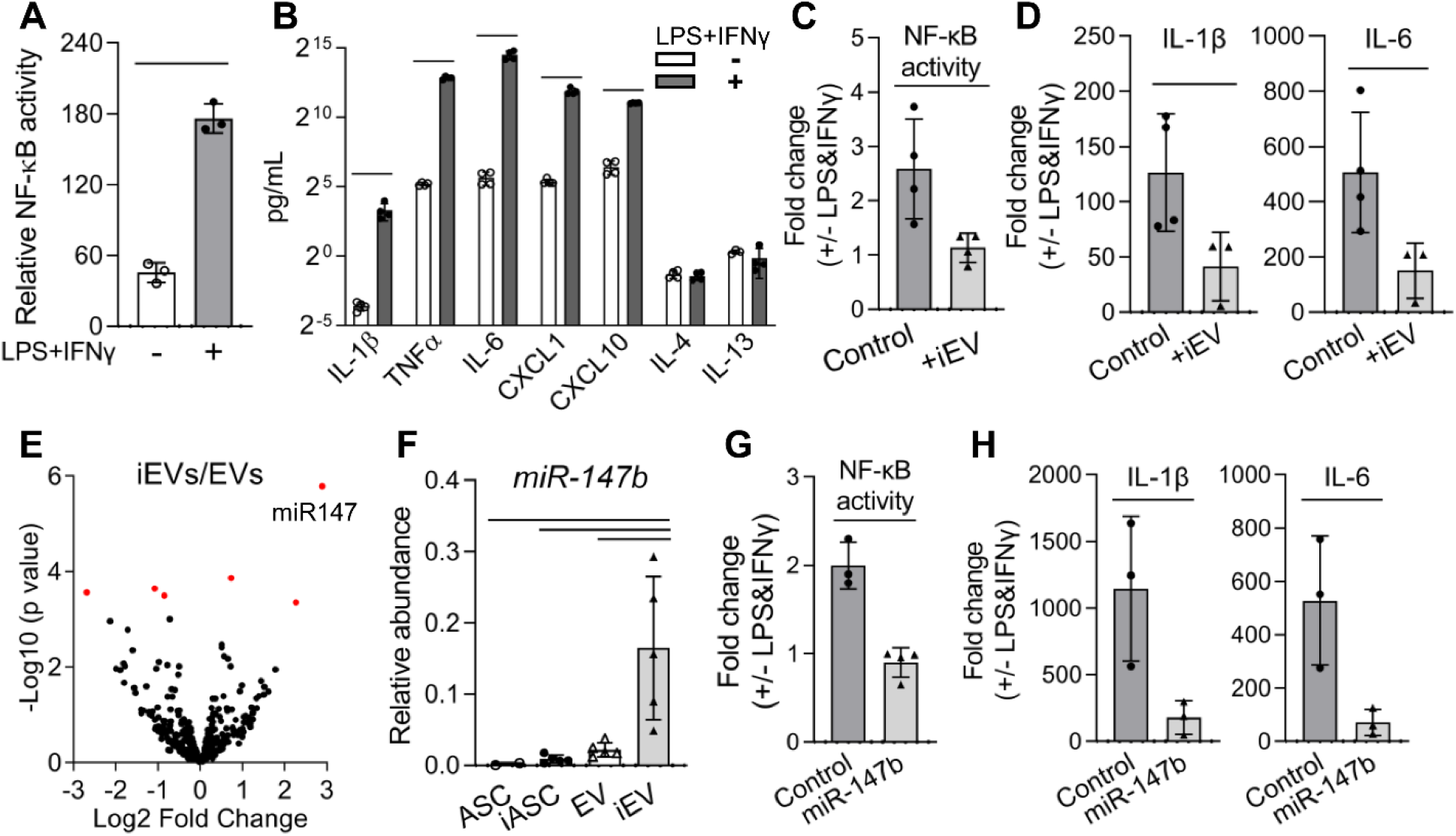
iEVs modulate macrophage inflammatory response by inhibiting toll-like receptor 4 (TLR4)/NF-κB signaling pathway with regulatory miRNAs. **(A)** Macrophage NF-κB activity induced by TLR4 agonists LPS and IFNγ. **(B)** Macrophage cytokine and chemokine release stimulated by LPS and IFNγ. **(C and D)** The effect of iEVs on macrophage NF-κB activity (C) and inflammatory cytokine release (D) induced by LPS and IFNγ. (**E**) Volcano plot of a small RNA-sequencing analysis showing differentially carried miRNAs between iEVs and EVs generated by inflammation-primed and naïve ASCs (iASC and ASC), respectively. • and •, adjusted p value < = 0.05 and > 0.05. **(F)** Relative abundances of miR-147b in iEVs, EVs, and their parent cells. **(G and H)** The effect of miR-147b mimic overexpression on macrophage NF-κB activity (G) and inflammatory cytokine release (H) induced by LPS and IFNγ. Control, control mimics. —, p < 0.05 between indicated groups for A-D and F-H.

To identify active molecules mediating the anti-inflammatory effect of iEVs, we performed a small RNA-sequencing analysis. Because iEVs have been found to be more effective than naïve EVs in reducing tendon inflammatory response,^22^ iEVs and their parent iASCs were compared with naïve EVs and ASCs. Results revealed that both naïve and primed EVs contained more miRNAs than parent cells (Supplemental Figure 1). Further differential analysis of iEV and naïve EV miRNAs identified miR-147 as the most distinctively enriched miRNAs in iEVs (p = 0.00072; Figure 6E). We validated the finding by miRNA Taqman PCR and confirmed that iEVs carried the most miR-147-3p (alias, miR-147b) compared to naïve EVs and their parent cells (Figure 6F). Overexpression of miR-147b but not negative control mimics in macrophages effectively blocked macrophage NF-κB activation (Figure 6G) and inflammatory cytokines IL-1β and IL-6 releases (Figure 6H) stimulated with LPS and IFN-γ.

### The in vitro effect of iEVs on tendon cells

Besides macrophages, iEVs have been found to target ScxGFP+ tendon cells in vivo.^22^ To better understand the effect of iEV on tendon cells, we co-cultured tendon cells and iEVs. Results confirmed that iEVs were taken up by tendon cells (Figure 7A), leading to a 33% increase in tendon cell population compared to untreated cells 48 hours after co-culture (Figure 7B). Significantly, the effect was coupled with a 45% increase in tendon cell type I collagen production after adjusted by the cell population increase (Figure 7C), thus demonstrating an anabolic effect of iEVs. Overexpression of miR-147b mimics in tendon cells didn’t recapitulate the effect of iEVs on tendon cell proliferation (Figure 7D, p=0.935), indicating other active molecules carried by iEVs mediated the effect.

**Figure 7.**
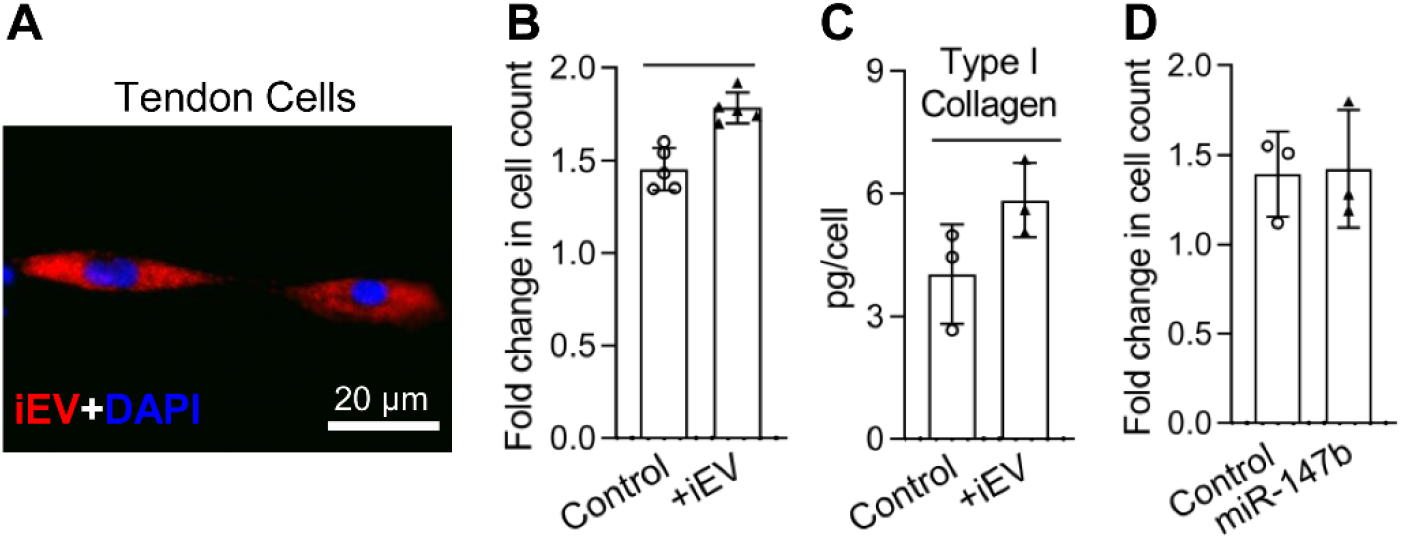
iEVs promote tendon cell proliferation and type I collagen production. **(A)** A representative fluorescence image of tendon cells stained with 4’,6-diamidino-2-phenylindole (DAPI) in blue and iEVs stained with PKH26 in red. **(B)** Fold change in tendon cell population 48 hours after vehicle control or iEV treatment. **(C)** Type I collagen produced by tendon cells 48 hours after vehicle control or iEV treatment. **(D)** Fold change in tendon cell population 48 hours after treatment with negative control miRNA mimics or miR-147b mimics. —, p < 0.05 between indicated groups for B and C.

## Discussion

A major clinical challenge for biological improvement of tendon repair is to achieve the contrasting goals of reducing inflammation without compromising tendon structure and strength recovery and stimulating tendon tissue regeneration while avoiding peritendinous scar formation. ^5,21,26,34,35^ Prior work has established the role of iEVs in attenuating inflammation in the early phase of tendon healing.^22^ It remained to be determined if iEVs might promote tendon structural and functional recovery while limiting scar tissue formation in the later phases of tendon healing. Here, we examined the long-term dose effect of iEVs in a clinically relevant mouse Achilles tendon injury and repair model. Results first confirmed that iEVs dose-dependently reduced repair site NF-κB activity in the early inflammation phases of tendon healing. Results further showed that iEVs increased anti-inflammatory gene expression and reduced inflammatory cell accumulation in the later remodeling phases of healing. Significantly, iEVs both promoted collagen deposition and tendon structural recovery and limited peritendinous scar formation, leading to less postoperative complications, and accelerated functional recovery. The subsequent in vitro studies further revealed that iEVs can reduce inflammation via miR-147b that targets macrophage TLR4/NF-κB pathway and promote tendon healing by increasing tendon cell proliferation and collagen production. Collectively, our results demonstrated that iEVs are a promising therapeutic agent for enhanced tendon repair by both reducing inflammation and stimulating intrinsic healing response.

Current results strongly support that the anti-inflammatory function of iEVs at least partially resulted from their ability to inhibit macrophage TLR4/NF-κB signaling by delivering anti-inflammatory miRNAs including miR-147b. Specifically, this and our prior study showed that iEVs effectively reduced *Tlr4* expression and NF-κB activation after Achilles tendon injury.^22^ iEVs were further found to block macrophage NF-κB activation and IL-1β and IL-6 production triggered by the TLR4 agonists LPS and IFNγ. The subsequent small RNA-sequencing study revealed that iEVs are distinctively enriched with miR-147b. Like iEVs, overexpressing miR-147b mimics inhibited macrophage NF-κB activation and IL-1β and IL-6 production induced by LPS and IFNγ. Consistently, miR-147 has been reported to reduce inflammatory cytokine expression in peritoneal macrophages stimulated with ligands to TLR4 in vitro^36^ and attenuate aortic inflammation as an active component of other stem cell extracellular vesicles in vivo.^37^

iEVs also targeted tendon cells and increased tendon cell proliferation and type I collagen production. Overexpression of miR-147b mimics in tendon cells showed no apparent effects on tendon cell proliferation, indicating other iEV molecules contributed to the effect. Besides miRNAs, iEVs carried other small non-coding RNAs and mRNAs, which may contribute to the iEV effects on tendon cells. Thus, further studies are needed to fully elucidate the cellular and molecular mechanisms of iEV action in enhancing tendon repair.

Our results showed dose-dependent effects of iEVs on tendon repair. As expected, high dose iEVs were more effective than low dose iEVs in reducing repair site NF-κB activity in the early inflammation phase of tendon healing and in promoting anti-inflammatory genes *Il1rn* and *Il13* expression and collagen deposition in the later remodeling phase of healing. Interestingly, in contrast to the enhanced tendon structural and functional recovery, iEVH-treated tendons expressed less *Col1a1* and *Scx* than untreated tendons in the remodeling phase. As tendon cell propagation and matrix production are expected to peak in the earlier proliferation phase of healing, the reduced tendon gene expression possibly reflects an accelerated healing process following iEV treatment. This idea is supported by our prior study, showing iEVs increased collagen deposition in the injury center by 3-fold as early as 7 days after repair.^22^ Moreover, excessive inflammation is known to cause disorganized collagen deposition and fibrotic scar formation, which impair tendon function.^34,35^ The reduced *Col1a1* expression in the remodeling phase of healing likely also a result of iEV’s ability to mitigate inflammation. Consistently, iEVs were found to effectively reduce mononuclear cell accumulation in repaired tendons, and the effect was coupled with reduced peritendinous scar formation.

Because Achilles tendon integrity is required for plantar flexion of the foot and such locomotion movement as running, the ankle joint angle of injured limb during running was used as an index for Achilles tendon functional recovery.^31^ As expected, the angle was significantly reduced following Achilles tendon transection. With less scar tissue formation and better tendon matrix and structural recovery, mice treated with either dose of iEVs exhibited higher recovery rates than untreated mice. While high dose iEVs more effectively reduced inflammation and promoted tendon matrix regeneration and remodeling than low dose iEVs, no apparent dose effect of iEVs on the recovery rate was detected. This discrepancy is likely because the running condition was set at an easy level (i.e., 6~8 m/min without slope), which limited the ability of this analysis to detect dose effect. Future assessment of tendon functional recovery will be conducted during uphill running at a higher speed (e.g., 12-16 m/min).

A limitation of this study is the lack of biomechanical analysis of repaired tendons. Although the ankle joint analysis demonstrated the ability of iEVs in alleviating mouse functional deficits after Achilles tendon injury and repair, further evaluation of the iEV effect on tendon biomechanical property could provide more useful information for clinical application of iEVs in treating tendon injury.

## Conclusion

Our results showed that iEVs can effectively enhance Achilles tendon healing by both targeting macrophages to attenuate inflammation and tendon cells to promote tendon tissue regeneration and functional recovery. Results also identified miR-147-3p as one of active components mediating the anti-inflammatory function of iEVs. These findings paved the road toward clinical application of iEVs in tendon repair and provided the basis for future engineering component-defined and disease-specific iEVs to treat tendon injury and other related disorders.

## Acknowledgement

This research was funded by the NIH/NIAMS R21AR075274, the Foundation for Barnes-Jewish Hospital, and the Washington University Institute of Clinical and Translational Sciences (supported by the NIH/NCATS UL1 TR002345). The authors thank Dr. Ratna B. Ray for her help with developing this project and reading the manuscript as well as Dr. Dimitrios Skouteris for his assistance with mouse surgeries. The authors also thank Crystal Idleburg and Samantha Coleman in the Washington University Musculoskeletal Research Center (supported by the NIH/NIAMS P30 AR074992) for preparing tendon sections. This publication was made possible in part by NIH/NCRR UL1 RR024992, which supported the RNA-sequencing service provided by the Washington University Genome Technology Access Center.

## Author Contributions

HS: Conception and design, funding acquisition, collection and assembly of data, data analysis and interpretation, manuscript writing, and final approval of manuscript

RAL: Collection and assembly of data, data analysis, and final approval of manuscript

