## Supplemental Figure 1 for "Extracellular Vesicles from Inflammation-Primed Adipose-Derived Stem Cells Enhance Achilles Tendon Repair by Reducing Inflammation and Promoting Intrinsic Healing"

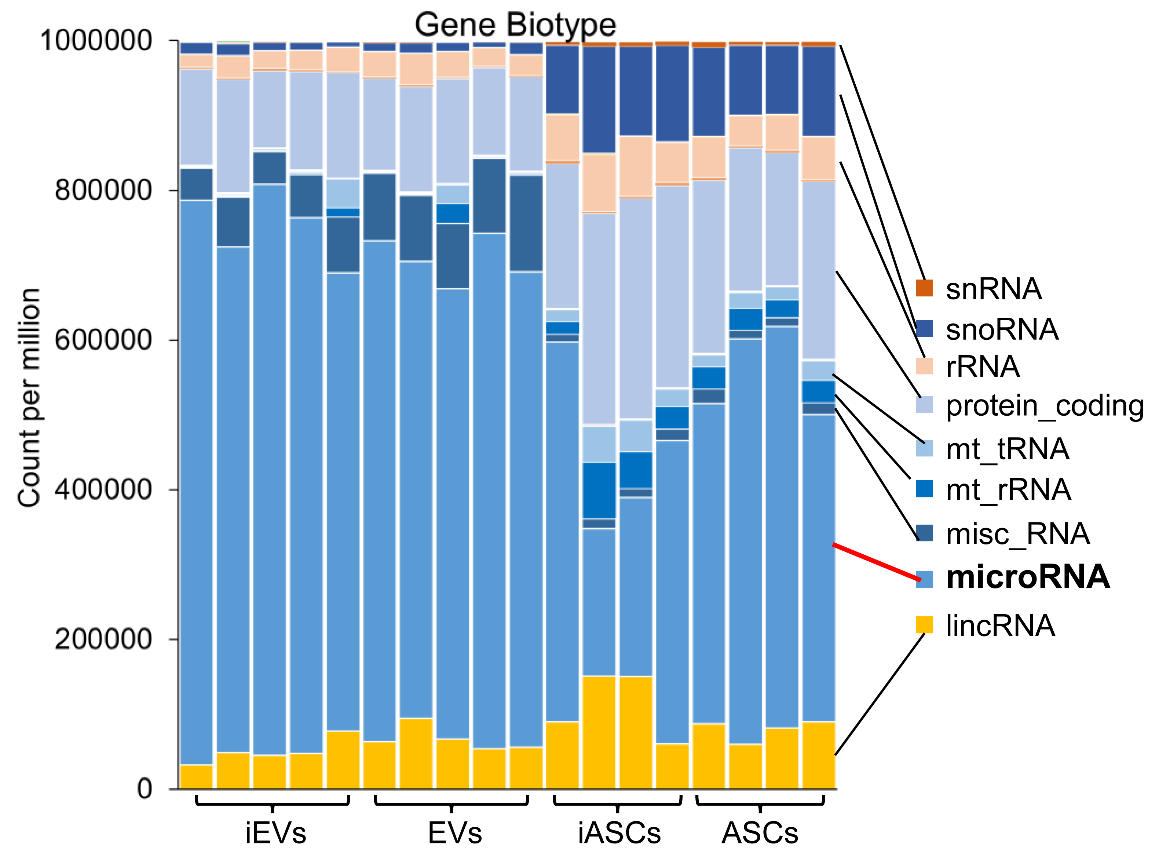
**Supplemental Material**

**Supplemental Figure 1.** Gene biotypes of extracellular vesicles and their parent cells. iEVs and EVs, extracellular vesicles from inflammation-primed and naïve adipose-derived stem cells (iASCs and ASCs).
